# Sperm Cell Painting: A Mechanism Driven Approach for Drug Discovery in Human Spermatozoa

**DOI:** 10.1101/2023.09.15.557919

**Authors:** Zoe C Johnston, Franz S Gruber, Rachel Myles, Shruti V. Kane, Anthony Richardson, David P. Day, Caroline Wilson, Kevin D. Read, Ian H. Gilbert, Sarah Martins da Silva, Christopher LR Barratt, Jason R. Swedlow

## Abstract

We have adapted the cell painting assay developed by Carpenter and colleagues on cultured U2OS cells to human spermatozoa. In Sperm Cell Painting (SCP) we assemble an image-based quantitative fingerprint of the functional state of sperm. We use this assay to gain insight into the mechanism of action of compounds that modify sperm function and as a platform for contraceptive discovery.

## MAIN TEXT

Despite recent advances in the tools and technologies used to study human reproductive health, and a resurgence in interest around nonhormonal contraception, there remains a lack of knowledge and understanding of the fundamental biology underpinning human sperm function. This gap in understanding limits the diagnosis and treatment of infertility and slows the development of effective new contraceptive options.

To begin to address these gaps in understanding, we have recently developed a high-throughput phenotypic screening platform in human sperm and used it to screen for compounds that either both decrease [1] and enhance [2] sperm motility. This platform presents opportunities for target-agnostic screening which can be adapted for screening of specific sperm functions (reviewed in Johnston *et al*., 2021 [3]). Target agnostic phenotypic assays provide a powerful, unbiased approach to drug discovery, but require substantial downstream follow-up of any hits to identify targets and assess the opportunity for further compound development. Each assay is also limited to the examination of individual sperm functions such as motility or acrosome reaction.

Cell Painting is an image-based, high-content method using multiplexed fluorescent dyes for cytological profiling [4]. Images of cells can be converted to quantitative feature fingerprints and be used in screening assays and to classify [5, 6] the mechanism of action (MOA) and/or toxicity of drugs and drug candidates, genetic perturbations[5, 6] (Figure 1A) and is particularly powerful when combined with curated libraries of references compounds with known MOA, or known phenotypic effect, such as the Sperm Toolbox [7].

**Figure 1.**
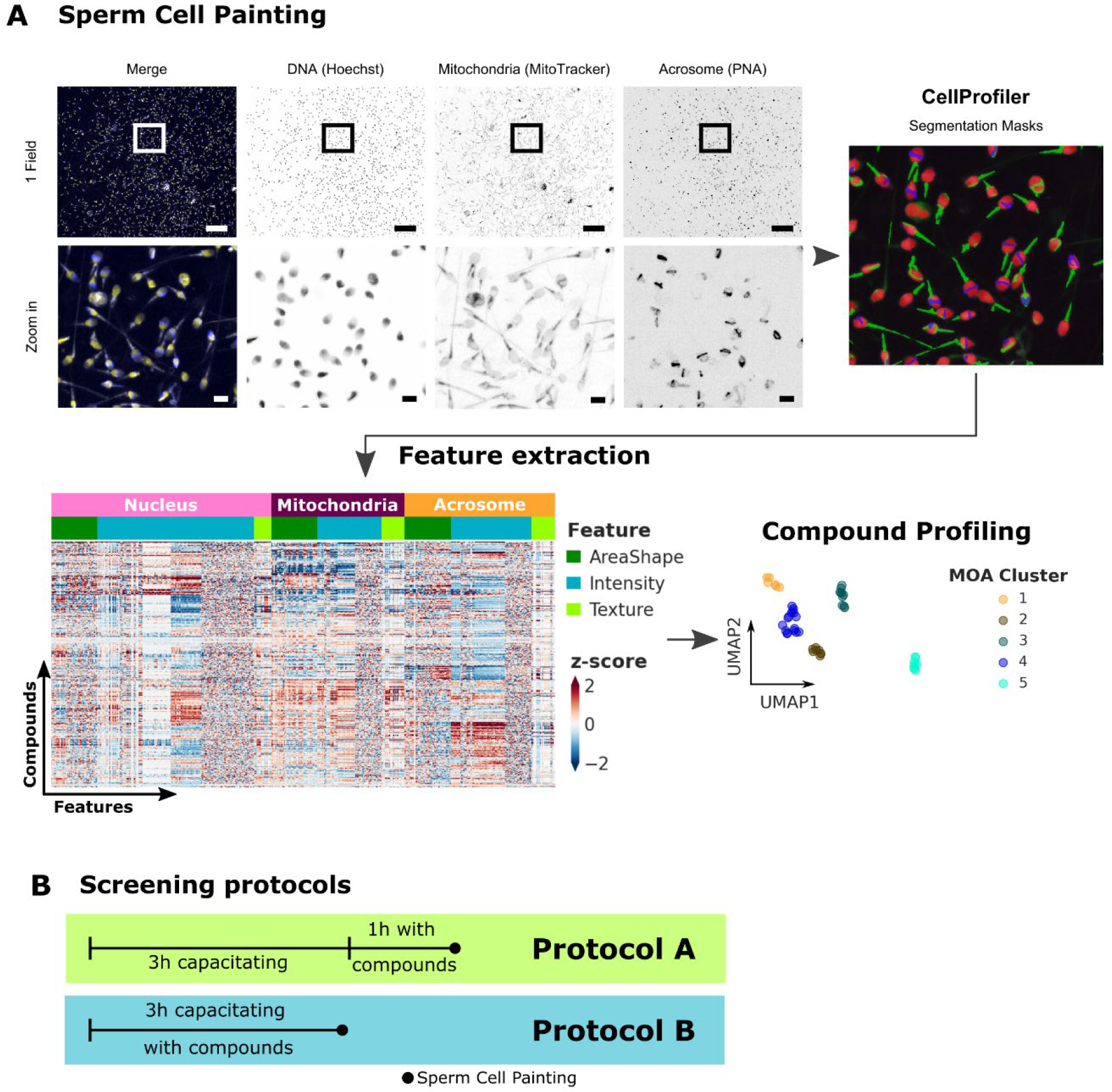
Sperm Cell Painting Overview (A) Sperm Cell Painting is based on the Cell painting protocol by Bray et al., 2016 [4]. After compound treatment, three sperm organelles are stained: Nuclei (Hoechst), Mitochondria (MitoTracker) and Acrosome (PNA, peanut agglutinin). Image segmentation and feature extraction is performed using CellProfiler. After data normalization, feature profiles of compounds can be compared and analysed for similarities to inform compound mode of action. Scale bar: Top row (1 Field) 50 μm, bottom row (Zoom in) 5 μm. (B) Schematic of the screening protocols used for exposure Protocol A and B. Exposure Protocol A: short exposure of capacitated cells. Cells were incubated in conditions which support capacitation for 3 hours followed by 1 hour of compound incubation in the same medium. Exposure Protocol B: long exposure of cells during capacitation. Cells were exposed to compounds during the 3 hours of incubation in conditions which support capacitation.

In this study, our aim was to adapt this powerful, high-throughput approach for use with human spermatozoa. Unlike the adherent, somatic cells that have previously been categorised by cell painting [4, 8], spermatozoa are not only motile, but lack some of the cellular components/machinery of somatic cells such as the endoplasmic reticulum. Therefore, significant adaptations are required to utilise the techniques with human sperm cells (see Methods).

A major goal of this work was to develop a phenotypic screening platform for the discovery of a female-controlled contraceptive that targets spermatozoa.. After ejaculation, spermatozoa experience different physiological states as they travel within the female reproductive tract. In response, they undergo a process known as capacitation, which involves destabilization of the acrosome membrane and changes in the tail that promote motility and fertilisation competence (reviewed in Puga Molina et al., 2018 [9]). Although the physiological and behavioural changes associated with capacitation have been studied in detail, the molecular basis for capacitation is not well-characterized. For contraceptive development, it is therefore an ideal functional target, but identification of molecular targets will require a screening strategy that provides as much mechanistic detail as possible. In addition, as the process of capacitation is highly species-specific, we focussed on assays that employed living human sperm to ensure compound hits would have effects in humans.

For these reasons, we have designed an assay strategy that supports capacitation in human sperm, and designed compound incubation protocols (Figure 1B) that model short (1 hour; Protocol A) compound exposure on capacitated cells and long exposure (3 hours; Protocol B) on cells during the capacitation process, allowing us to model the effects of compound exposure at different points during the journey of sperm through the female reproductive tract.

We incubated compounds from several small, well annotated libraries with living spermatozoa and then processed cells for Cell Painting (Figure 2A). We used CellProfiler to identify cells and segment nuclei, acrosome membrane and the mitochondria and calculate image-based features. Feature normalisation and scaling was performed as described in Bray et al (2016; [4]). We filtered compounds using an induction score (Figure 2B), and selected compounds for further analysis where >15% of features were >= 2σ deviation from DMSO (Figure 2C), which produced 140 and 146 compounds using Protocol A and B respectively (Figure 2D), of which 56 were observed in both exposure protocols (Figure 2E). The pattern of z-scores in the heatmaps of the two are visibly different (Figure 2 F & G), indicating differential responses to compounds by cells subject to different exposure protocols. Furthermore, we observed that the annotated mechanism of actions of these compounds are similar, but different with different compound exposure protocols (Figure 2H). Both exposure protocols share a majority of mechanisms of action, however Protocol A detects more estrogen and dopamine receptor related compounds, while Protocol B detects sodium channel, serotonin receptor and cyclooxygenase related compounds. This highlights the potential for information gained from this approach to define pathways for further screening or for target deconvolution, and the importance of selecting screening conditions that reflect the relevant physiological state of the target cell.

**Figure 2.**
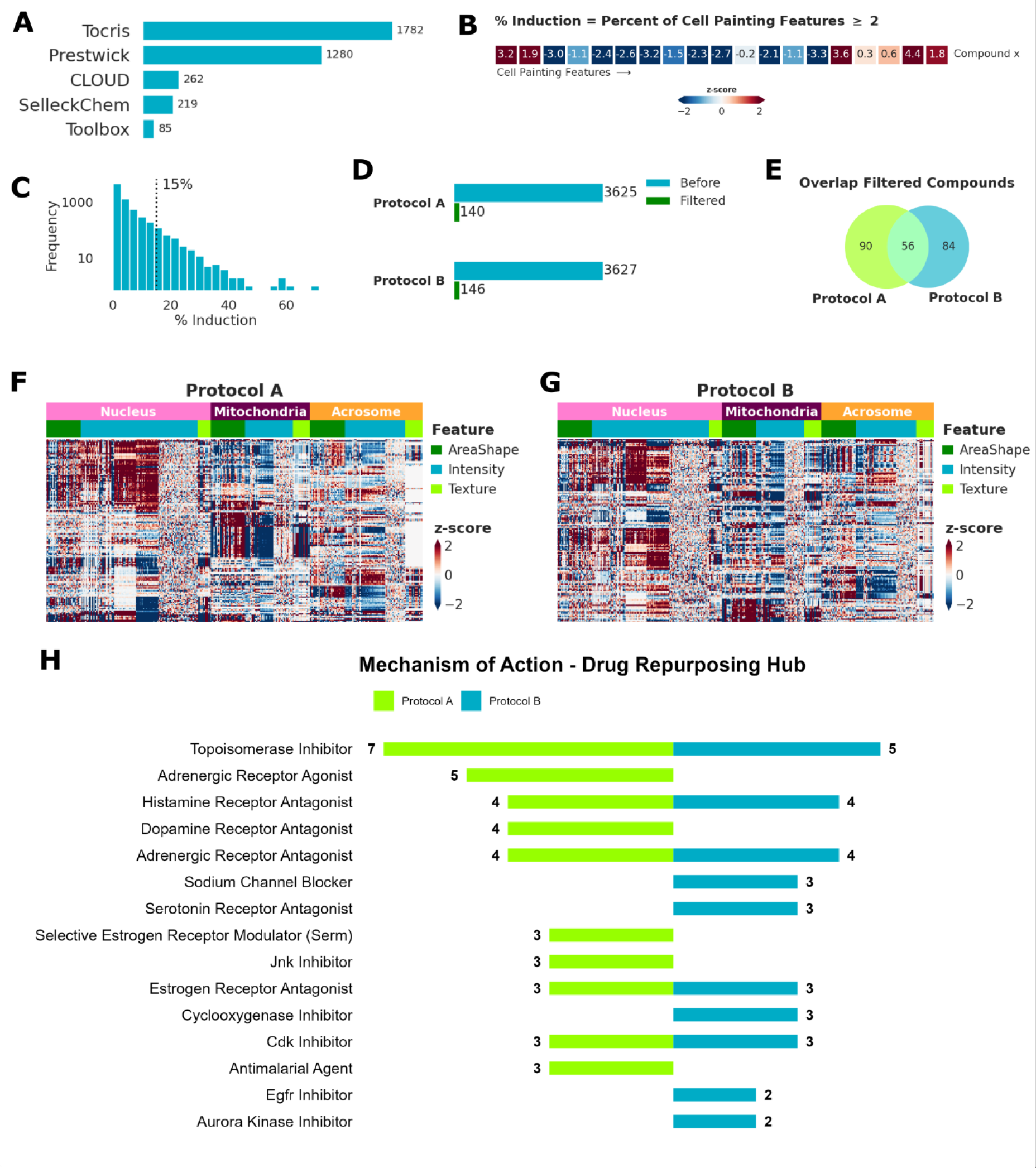
Summary of compound screening (A) Summary chart of screened libraries in this study. (B) Definition of ss Induction as the percentage of cell painting features greater than or equal to 2. (C) Frequency distribution of % Induction of all compounds screened. Dotted line indicates 15% cut-off for compound filtering. (D) Bar chart before and after filtering compounds using 15 % Induction cut-off. (E) Overlap of filtered compounds comparing Protocol A and Protocol B. (F) Heatmap showing z-scores (Red-White-Blue) of filtered compounds (rows) and features (columns) after **Exposure Protocol A**. Colors indicate organelles Nucleus (pink), Mitochondria (purple), Acrosome (orange) or feature type AreaShape (dark green), Intensity (blue), Texture (light green). (G) Heatmap showing z-scores (Red-White-Blue) of filtered compounds (rows) and features (columns) after **Exposure Protocol B**. Colors indicate organelles Nucleus (pink), Mitochondria (purple), Acrosome (orange) or feature type AreaShape (dark green), Intensity (blue), Texture (light green). (H) Summary of top 15 annotated mechanism of action for exposure Protocol A (light green) and exposure Protocol B (blue)

To better understand similarities between compounds and exposure protocols, we calculated pairwise correlation distance and generated cluster labels using hierarchical clustering (Figure 3A). This shows that there are multiple mechanistic clusters sharing profile similarity. In addition, we used UMAP dimensionality reduction to visualise groups of compounds with similar feature profiles to each other in the Sperm Cell Painting assay (Figure 3B). As with the heatmap, UMAP revealed that both incubation states generate similar overall patterns, but with visible differences (Figure 3C). This result confirms our hypothesis that different target engagement occurs in cells in different physiological states or with different incubation times.

**Figure 3.**
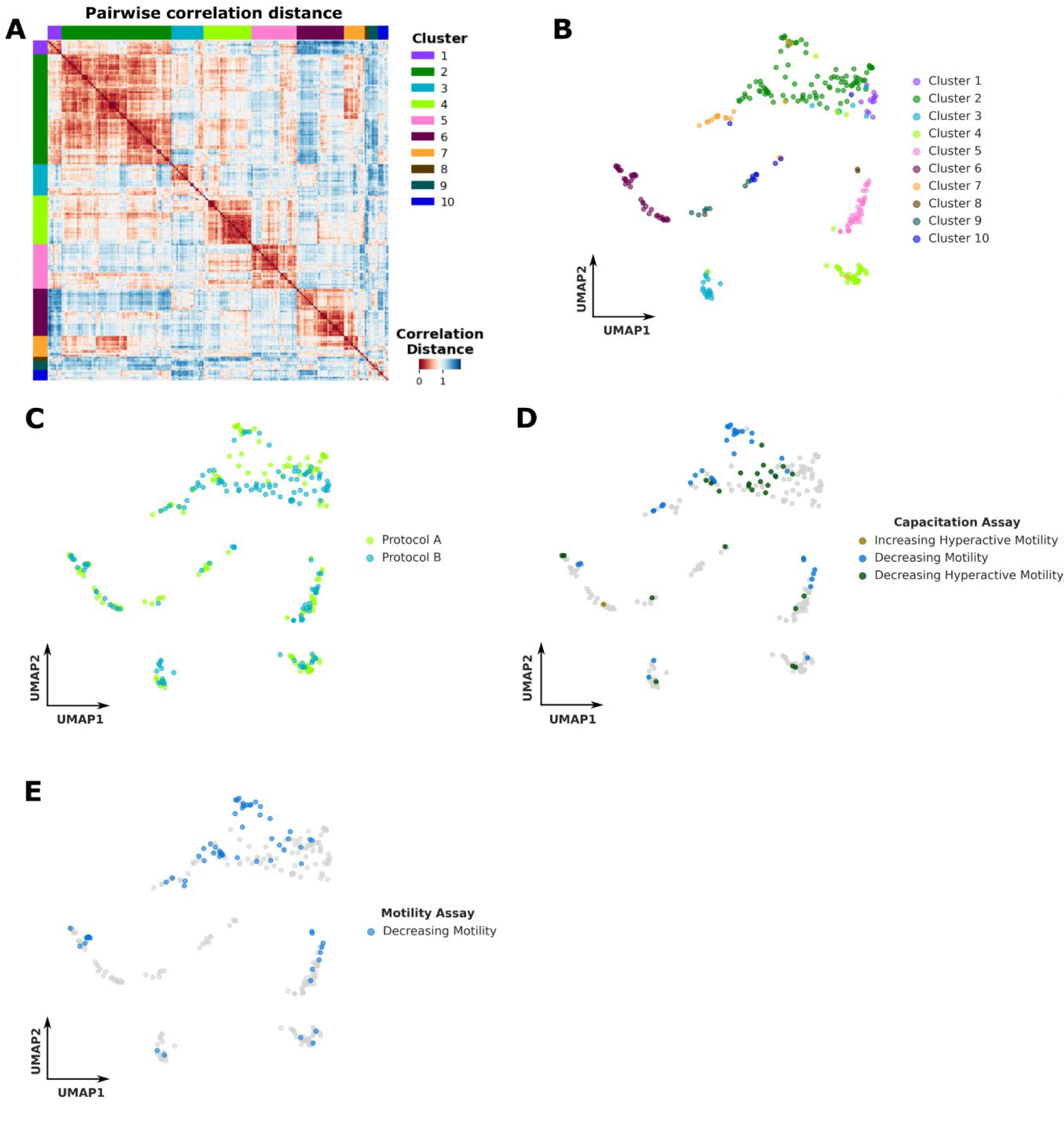
Feature profile comparison (A) Pairwise correlation distance calculation of compounds and hierarchical clustering of all filtered compounds (>15% induction). Color indicates profile similarity: red (more similar) blue (more distinct). Row and column color indicates cluster label. (B) UMAP visualization of filtered compounds (>15% induction). Color indicates hierarchical cluster label (same as in (A)). (C) Mapping compound exposure protocol onto the UMAP visualization. Color indicates exposure Protocol A (light green) or Exposure Protocol B (blue) (D) Mapping capacitation assay results onto the UMAP visualization. Color indicates compound effect in this assay. (E) Mapping motility assay results onto the UMAP visualization. Color indicates whether a compound shows >70% reduction in progressive motility in dose response experiments.

These visualisations can also be used to map phenotypes in orthogonal sperm assays. Compounds which either increase or decrease hyperactive motility in our capacitation assay (See online methods – Motility Assays) are dispersed throughout each of the clusters (Figure 3D). These effects are seen both in addition to, and independently from changes to progressive motility for individual compounds (Figure 3D). Similarly, cross-referencing these clusters with datasets from our original non-capacitated motility assay (See online methods – Motility Assays) reveals that hits from the motility and capacitation assay do not lie within any one cluster (Figure 3E). These results are consistent with the hypothesis that multiple molecular pathways are likely to be involved in the regulation of motility and capacitation [9].

One potential application of the Sperm Cell Painting assay is to look for compounds which affect the acrosome reaction (AR). We have used control compounds in our assays which artificially induce AR, such as A23187 a well characterized Ca2+ ionophore. A23187 is part of Cluster 2 (Figure 4A). This cluster contains an accumulation of compounds that have an annotated mechanism of action related to adrenergic, histaminergic, and serotonergic receptors (Figure 4B). Images of compounds with a similar feature profile as A23187 show that these compounds induce very high levels of typical acrosomal banding pattern staining, suggesting that they do induce acrosome reaction (Figure 4C). These data suggest that targets related to serotonin receptors, a receptor coupled GTPase, and at least one protein kinase are potential, previously unknown, targets for premature induction of AR and thus candidates for drug development programs. Combining Sperm Cell Painting with target-directed assays will be a powerful approach for future contraceptive and fertility drug development. The induction of the acrosome reaction at high frequency also confirms that our assay are effectively supporting capacitation.

**Figure 4.**
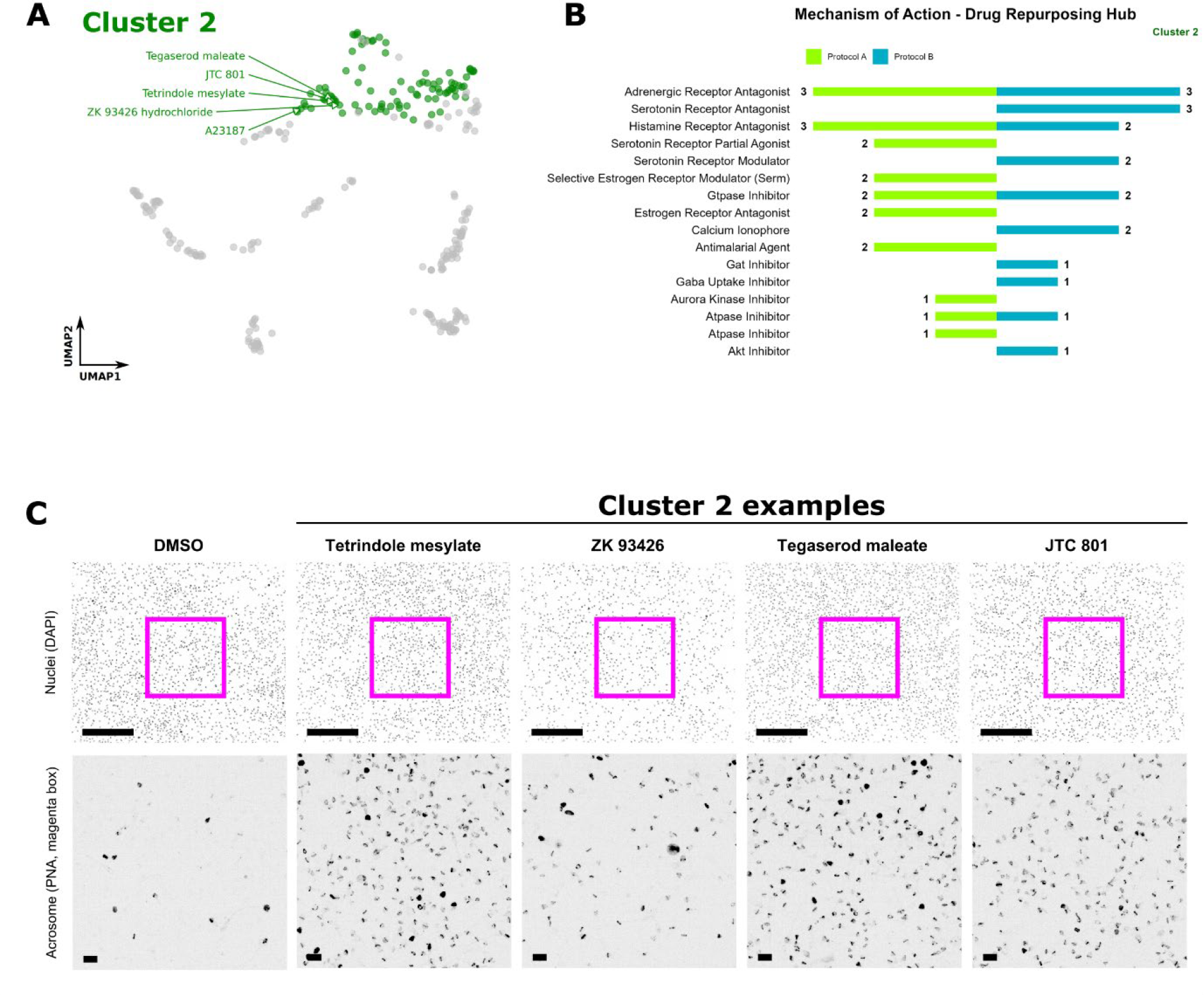
Cluster 2 contains compounds inducing acrosome reaction (A) UMAP of filtered compounds indicating compounds in cluster 2 (green labels), which share a similar profile as A23187 (acrosome inducing Ca2+ Ionophore). (B) Summary of the top 10 mechanism of action of compounds in Cluster 2. Compound annotation (MOA) used from drug repurposing hub. Colors represent exposure modes A (light green) and B (blue). (C) Visual confirmation of compounds indicated in (A). Top row showing nuclear staining (Hoechst), bottom row showing acrosome staining (PNA). Magenta box indicates zoom in region. Scale bars, 100 μm top row, 10 μm bottom row.

In summary, we have successfully transferred the well-established Cell Painting assay to a non-somatic and motile cell type. We observe differences related to the exposure Protocol used, highlighting the importance of screening in conditions which reflect the appropriate physiological state of the cell for the target application. We also observe mechanistic clusters of compounds sharing similar feature profiles, which presents an advantage for efficient drug discovery programmes. The ability to combine mechanistic and phenotypic approaches to screen and compare the data from the Sperm Cell Painting assay with that from other phenotypic assays has enabled us to create rich datasets that could prove invaluable in driving forward knowledge in male (in)fertility and drug-discovery. The use of well-defined and annotated libraries such as the Sperm toolbox [7] further increases the relevance of the information gained from screens to benchmark new assay and define areas of human sperm biology.

## ONLINE METHODS

### Experimental Design

We developed and utilised a cell painting platform for human spermatozoa. The cell painting technique was adapted for staining and fixation of non-adherent and motile cells, and used in conjunction with our automation and high-throughput screening platform to screen 3,628 compounds for their effects on sperm cell features. Custom CellProfiler pipelines were utilised for data analysis and percentage induction of selected features was used to determine hit compound.

### Spermatozoa

Semen samples were obtained from volunteer donors by masturbation, after a sexual abstinence period of 2-5 days. Written consent was obtained from each donor in accordance with the Human Fertilization and Embryology Authority (HFEA) Code of Practice (version 8) under local ethical approval (SMED REC 20/45). Donors were selected based on normal semen characteristics, sperm concentration, and motility according to WHO criteria (2010).

#### Media

##### Non-capacitating media (NCM)

Non-capacitation media (Minimal Essential Medium Eagle Sigma M3024, supplemented with 25mM HEPES (Invitrogen 15630056), 41.75 mM Sodium DL-lactate (Sigma L4263), 3.75mM Sodium pyruvate (ThermoFisher Scientific 11360070) and 0.3% Bovine Serum Albumin (Roche BSAV-RO) to achieve concentrations previously described [10].

##### Human Tubal Fluid/Capacitating media (HTF)

Capacitating media was composed of 97.8 mM NaCl (Sigma 230391), 4.69 mM KCl (Sigma P5405), 0.2 mM MgSO4 (Sigma 230391), 0.37 mM KH2PO4 (Sigma P0662), 2.04 mM CaCl2(Sigma 21115), 0.33 mM Na-pyruvate, 21.4 mM Na-lactate, 2.78 mM glucose (Sigma G8270), 21 mM HEPES, 25 mM NaHCO3 (Sigma S5761) and 0.3% Bovine Serum Albumin, adjusted to pH 7.4 with NaOH.

Samples were allowed to liquify at 37°C for 15 to 30 minutes prior to preparation. Spermatozoa were isolated from semen samples by discontinuous density gradient centrifugation (DGC), using 80% and 40% Percoll (Sigma Aldrich, UK). Gradients were prepared with non-capacitation media (NCM).

To reduce screening batch variability, 3-5 prepared donor samples were pooled to create each screening batch. Once pooled, samples were resuspended in HTF to a concentration of 5 × 10^6^/mL and incubated at 37°C, in 5% CO_2_ for either 30 minutes (protocol B) or 3 hours (protocol A).

### Sperm Cell Painting - High-throughput screening

#### Assay ready plates

Assay-ready 384-well plates were prepared prior to the screen. Library compounds (10mM stocks in DMSO) were dispensed using an acoustic dispenser (Echo 555, Labcyte Inc) to the desired concentration in poly-D-lysine-coated plates (poly-D-lysine-coated PhenoPlate 384-well; PerkinElmer; 6057500). The positions of each compound in the plate were allocated at random, and DMSO concentration was consistent between wells.

#### Libraries screened

From the Tocris compound library, we screened 1783 biologically active compounds selected from Tocriscreen plus (Tocris, Bristol, UK, https://www.tocris.com/products/tocriscreen-plus_5840) and Tocriscreen 2.0 (https://www.tocris.com/products/tocriscreen-2-0-max_7150).

The Prestwick Chemical Library contains off patent drugs with known human bio-availability and safety profiles (https://www.prestwickchemical.com/screening-libraries/prestwick-chemical-library/) and contained 1280 compounds at the time of purchase.

The CeMM library of unique drugs (CLOUD; https://enamine.net/compound-libraries/bioactive-libraries/the-comprehensive-drug-collection-cloud) contained 262 compounds at the time of purchase. The library contains a set of small molecules representing the chemical space of FDA-approved drugs.

The SelleckChem library contains 219 cherry picked annotated bioactive compounds.

The Sperm Toolbox is a library of 85 small molecules assembled specifically for the study of human spermatozoa [7].

#### Compound exposure

Two compound exposure protocols were used in order to examine the effects of compound exposure on cells in different physiological states. For both cases, screening batches of cells were transferred to a robotic platform (HighRes Biosolutions Inc.) and maintained at 37°C. Cells were dispensed into assay-ready 384-well plates at approximately 100,000 spermatozoa per well using a using a liquid handling system (MultiDrop Combi; ThermoFisher). Cells were then incubated in these assay plates, in the presence of compounds, for either 1 or 3 hours (Figure 1B).

##### Exposure Protocol A

Prepared sperm cells were incubated in conditions which support capacitation (HTF) for three hours, prior to being dispensed into assay-ready 384-well plates containing compounds. The cells, which had been given time to undergo capacitation prior to addition to the plates, were then incubated with compounds within these plates for 1 hour prior to staining and fixation.

##### Exposure Protocol B

After a 30 minute rest period, prepared sperm cells were dispensed into assay-ready 384-well plates containing HTF and compounds. Cells were then incubated for 3 hours within the assay plate, and were therefore exposed to compounds during capacitation.

#### Staining protocol

Plates were stained with a combination of dyes targeting the nucleus, acrosome, and mitochondria of the sperm cells. These dyes were Hoechst 33342 (Thermo Scientific; 62249), PNA 647 (Invitrogen; Lectin PNA from Arachis hypogaea [peanut], Alexa Fluor™ 647 Conjugate; L32460), and MitoTracker green FM (Invitrogen; M7514) respectively.

An intermediate staining stock solution of the three dyes was diluted in HTF (10x final concentration).

A TEMPEST® liquid dispenser (FORMULATRIX) was used to dispense 2μL of the intermediate staining stock solution into each well of the assay plate, to achieve a final dilution of 1μL per 1mL Hoeschst 33342, 0.5μL per 1mL PNA 647, and 0.1μL per 1mL MitoTracker green FM. Plates were incubated for 30 minutes at 37°C

#### Fixation and adherence

Stocks of 4% PFA were prepared from PFA powder (Sigma 441244) in MQ-H2O and pH adjusted to pH 7 with addition of NaOH. An equal volume (20μL) of 4% PFA was dispensed by TEMPEST to each well, for a final concentration of 2% PFA, and incubated for a further 10 minutes at room temperature. To adhere the now-fixed cells to the poly-D-lysine-coated well bottom, each plate was centrifuged at 300g for 5 minutes in each of two orientations. The microplates were then washed gently using an automated Bio Tek 405 LS Washer (Agilent)

#### Imaging

Images were captured on an ImageXpress Micro XLS epifluorescence microscope (Molecular Devices) or CV7000 (Yokogawa) in three fluorescent channels.

### Data handling and analysis

#### Data Storage and processing

Data is stored in OMERO [11], along with metadata annotations and segmentation masks generated by CellProfiler. Images are processed using custom CellProfiler pipelines to segment nuclei, mitochondria and acrosomes. Segmentation masks are used to calculate AreaShape, Intensity and Texture measurements, which are used for downstream analysis.

#### Data normalisation

Data was normalized using a strategy adapted from Bray et al., 2016. In short, for each feature we calculate DMSO median and median absolute deviation (MAD), which are used to standardize (subtract median, divide by MAD) each feature to unit norm at the cell level. This is done plate wise, where each plate contains 16 wells of DMSO treated cells. Next, we calculate a well median of each feature. Finally, we perform a plate wise standardization using the mean and standard deviation to generate z-scores for each of the features. Data normalization is done using custom python scripts utilizing the power of pandas and numpy.

#### Feature selection

We remove highly correlated features (Pearson correlation > 0.98) and features with low variance (threshold 0.05%). Feature selection is done using custom python scripts utilizing pandas and scikit-learn (VarianceThreshold).

#### Percent induction

For each compound we calculate % percent induction as the percentage of features that have an absolute z-score of ≥ 2, similar as done by Akbarzadeh et al., 2021. We used an arbitrary cut-off of 15% induction to remove compounds with weak profiles.

### Hit analysis

Filtered compounds (≥ 15% induction) have been selected for further analysis. Pairwise correlation distance was calculated using SciPy (pdist, method = ‘correlation’). Hierarchical clustering was performed on the correlation distance matrix using SciPy (average linking) from which ten clusters were formed using SciPy (fcluster).

We used UMAP (McInnis et al.) as an orthogonal approach to reduce dimensions and visualize similarities among compound profiles.

#### Subcluster analysis

We augmented our dataset using curated data from the drug repurposing hub (https://clue.io/repurposing) to infer mode of action (MoA). In addition, we used the Bioconductor package clusterprofiler to study gene ontology (GO) enrichments or potential protein targets.

Throughout the paper data was visualized using python (matplotlib, seaborn) or R (ggplot2, clusterprofiler)

### Motility Assays

Compound induction profiles in the sperm cell painting assay were compared to the effects of compound on sperm motility using two forms of high-throughput motility assay.

Progressive motility was measured using the high-throughput assay previously published in Gruber *et al*., 2020 and 2022 [1, 2] in non-capacitating conditions and Non-capacitation media.

The effect of compound exposure on the hyperactivated motility of sperm cells was measured using a modification of this assay platform that we have termed the capacitation assay. Prepared cells were incubated in HTF medium for 30 minutes at 37°C and 5% CO_2_ prior to dispensing into an un-coated assay ready plate. Plates were further incubated for 30 minutes at 37°C and 5% CO_2._ After 30 minutes compound exposure, plates were imaged using the CV7000 (Yokogawa) and cell motility tracked and classified as in Gruber *et al*., 2020 [1]. Hyperactivated motility was defined using fractal analysis with VCL >= 150 um/sec and D>=1.2, adapted from Mortimer et al., 1991 [12].

## Data Availability

We will make these datasets available to the community through publication on IDR.

## DECLARATIONS

### Conflicts of interest

CLRB is Editor for RBMO. No other authors declare a conflict of interest.

### Authors Roles

ZCJ and FSG contributed equally to this paper. ZCJ developed and optimised the sample preparation, fixation and adhesion, and staining protocol. ZCJ and RM recruited donors and carried out cell preparation. ZCJ, RM, and FSG carried out compound screening. FSG developed the data analysis pipelines and FSG and SVK carried out data analysis and prepared data for inclusion in IDR. CLRB, KDR, IHG, SMDS, and JRS obtained funding and contributed to experimental design and data interpretation. AR, DPD, and CW contributed chemistry and compound handling expertise. ZCJ and FGS prepared the first draft of the manuscript. All authors contributed to subsequent drafts and to the final editing of the manuscript. All authors approved submission of the final manuscript.

### Funding

This work was supported by the Bill & Melinda Gates Foundation (INV-007117)

## Notes

### Competing Interest Statement

The authors have declared no competing interest.

### Summary of Updates

Author List Updated

